# Biased parasitoid sex ratios: *Wolbachia*, functional traits, local and landscape effects

**DOI:** 10.1101/271395

**Authors:** Zoltán László, Avar-Lehel Dénes, Lajos Király, Béla Tóthmérész

**Author notes:** Zoltán László (Corresponding author): E-mail address.

## Abstract

Adult sex ratio (ASR) is a demographic key parameter, being essential for the survival and dynamics of a species populations. Biased ASR are adaptations to the environment on different scales, resulted by different mechanisms as inbreeding, mating behaviour, resource limitations, endosymbionts such as *Wolbachia*, and changes in density or spatial distribution. Parasitoid ASRs are also known to be strongly biased. But less information is available on large scale variable effects such as landscape composition or fragmentation. We aimed to study whether the landscape scale does affect the ASR of parasitoids belonging to the same tritrophic gall inducer community. We examined effects of characteristics on different scales as functional trait, local and landscape scale environment on parasitoid ASR. On species level ovipositor length, on local scale resource amount and density, while on landscape scale habitat amount, land use and landscape history were the examined explanatory variables. We controlled for the incidence and prevalence of *Wolbachia* infections. Parasitoid ASR is best explained by ovipositor length: with which increase ASR also increases; and available resource amount: with the gall diameter increase ASR decreases. On large scale the interaction of functional traits with habitat size also explained significantly the parasitoid ASRs. Our results support the hypothesis that large scale environmental characteristics affect parasitoid ASRs besides intrinsic and local characteristics.

## Introduction

The adult sex ratio (ASR, ratio of adult males to females in a population) has critical effects on the ecology and population dynamics of insects and animals (Pipoly, Bókony, Kirkpatrick, Donald, Székely, et al., 2015). It is expected to be 1:1, which is the equilibrium ratio. Reasons for this are known as the Fisher’s principle (Fisher, 1930). However, ASR ranges from populations that are heavily male-biased to those composed only of adult females (Xu, Fang, Yang, Dick, Song, et al., 2016). Identification of causes and consequences of this variation has an extreme importance in population biology and biodiversity conservation because it affects the fitness of populations through breeding systems (Pipoly et al., 2015).

Biased ASRs may be adaptations to conditions on different scales, such as inbreeding due to small population sizes, resource limitations, changes in density or spatial distribution (Kraft & Van Nouhuys, 2013). Different factors leading to biased ASRs may be populational as sex-differential mortalities of young and adults, sex-differential dispersal and migration patterns (Székely, Liker, Freckleton, Fichtel, & Kappeler, 2014). On an infra-individual level, biased ASRs may be caused by reproductive parasites (endosymbionts) such as *Wolbachia* or *Cardinium* in many arthropod species (Floate & Kyei-Poku, 2013) as they kill males (Werren, 1997) and cause parthenogenesis (Provencher, Morse, Weeks, & Normak, 2005; Duplouy, Couchoux, Hanski, & Van Nouhuys, 2015).

Several related populational factors may alter parasitoid ASR such as female wasp density and host density (King, 1987). The local resource competition (LRC) theory (Clark, 1978) explains male biased ASR with the reduction of competition between daughters with small dispersal distances for local limiting resources (West, 2009). Local mate competition (LMC) (Hamilton, 1967) occurs when male relatives with low dispersal abilities compete for mating opportunities, favouring female-biased sex allocation (Rodrigues & Gardner, 2015), because they are in higher number than females: for example in case of fig wasps (Herre, 1985).

Parasitoid ASRs are known to be mostly female biased (Hamilton, 1967; Charnov, Hartogh, Jones, & Assem, 1981). Egg laying females control their offspring’s sex as a function of host size. Haplodiploid sex determination provides parasitoid females a physiological mechanism for this control (Charnov et al., 1981). A population level mechanism is based on the prediction that the rarer sex in a population may have higher fitness, i.e. isolated females produce primarily daughters (Frank, 1986). As the number of females increases, the number of sons has to increase as they become rarer (King, 1987). Another population level mechanism is based on host density: at low host density brood size and sex ratio are strongly positively correlated, while at high density there is no such relationship (Kraft et al., 2013).

Functional traits as ovipositor length of parasitoids are also adaptations to suboptimal conditions which may also have significant effect on ASR as well (Sivinski, Vulinec, & Aluja, 2001; Sivinski & Aluja, 2001). For species with short ovipositors, hosts finding is difficult and therefore they may show low population densities and aggregated distributions (Alvarenga, Dias, Stuhl, & Sivinski, 2016). Species with low population sizes and aggregated distributions are more likely to avoid LMC by female biased ASRs (Alvarenga et al., 2016), while species with large population sizes are inclined to show male biased (West, 2009) or 1:1 ASRs.

Large scale effects on parasitoid ASR are virtually unknown. Available studies from this perspective target usually vertebrates (Amos, Balasubramaniam, Grootendorst, Harrisson, Lill, et al., 2013; Amos, Harrisson, Radford, White, Newell, et al., 2014; Reid & Peery, 2014). The scale of known or debated causes of biased ASR in parasitoids (for a review see (King, 1987)) are usually infra-individual (*Wolbachia* presence, genetic variability) or local (population size effects). Therefore, we aimed to analyse beyond known affecting variables also large scale patterns on parasitoid ASR as landscape composition, configuration and landscape history. Our study hypothesis is that biased parasitoid ASR are also affected by large scale variables beyond infra-individual and local ones. Our predictions were as follows: (i) a functional trait, the ovipositor length is positively related to parasitoid ASR: longer ovipositors may be adaptations to limited resources; (ii) a small scale variable, the available resource amount is negatively related to parasitoid ASR: since limited resources cause increase of female bias; (iii) large scale, e.g. landscape characteristics are indirectly related to the parasitoid ASR: since habitat amount changes are related to changes in available resource amount.

## Material and Methods

### Location and studied species

Data were collected in three consecutive years (2004-2006) on seven landscapes (Fig. 1) positioned on a South-East – North-West axis of 328 km through the Transylvanian Plateau (Romania) and the Great Hungarian Lowland (Eastern Hungary). Galls were collected from randomly chosen 50×50 m area plots (N=65) from habitats of the Robin’s pincushion or rose bedeguar gall (*Diplolepis rosae*). Plot locations within the sites varied between the measurement years, thus, each plot was sampled only once. Galls from each infected bush from all plots were collected in February and March of each year. We stored collected galls individually in plastic cups under standard laboratory conditions. Emerged specimens were separated, then preserved in 70% ethanol for identification. We counted the emerged female and male individuals separately for all analysed species.

**Fig. 1.**
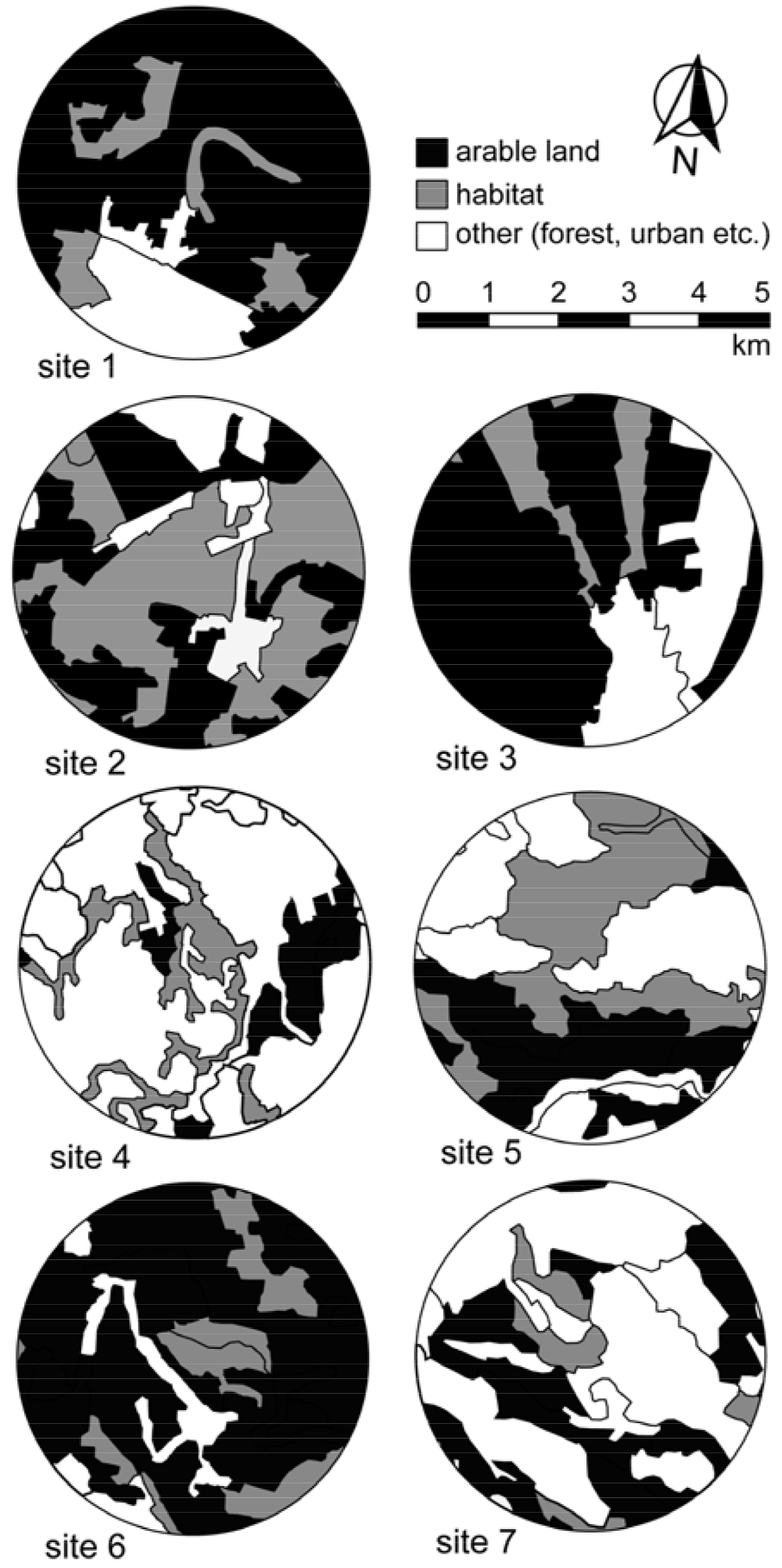
Collecting sites. The sampled landscape areas with habitat areas of the gall wasps and its parasitoids (bushy grasslands and pastures with shrub encroachments) and agricultural patches. Other patch types as forests, orchards, marshes and urban areas were not considered. Maps were acquired from Corine Land Cover 2006 vector layers.

The Robin’s pincushion induced by females of *D. rosae* has a Holarctic distribution, and is one of the most abundant cynipid galls in the Carpathian Basin and Eastern Europe. Gall wasp females produce multi-chambered galls on wild rose species without demonstrable preference for certain rose species (Kohnen, Wissemann, & Brandl, 2011). The most abundant primary solitary specialist parasitoid species of the *D. rosae* gall community in the Carpathian Basin are *Orthopelma mediator* (HYM: Ichneumonidae), *Pteromalus bedeguaris* (HYM: Pteromalidae), *Torymus bedeguaris* (HYM: Torymidae) and *Glyphomerus stigma* (HYM: Torymidae) (László, Rákosy, & Tóthmérész, 2014). We chose these species to analyse different scale effects on parasitoid ASR. *O. mediator* and *P. bedeguaris* emerge early in the spring (early flying species), when galls are small and have just begun to grow. *T. bedeguaris* and *G. stigma* emerge late in the spring (late flying species), when galls are large and close to maturation (László & Tóthmérész, 2011). Also, late flying species have significantly longer ovipositor sheaths than early flying species: ovipositor sheaths of *T. bedeguaris* and *G. stigma* are at least as long as combined length of meso-and metasoma, while for *O. mediator* and *P. bedeguaris* these are at most as long as metasoma (see Fig. 2 species habitus or keys of Gauld and Mitchell (1977), Graham (1969), and (Graham & Gijswijt, 1998)).

**Fig. 2.**
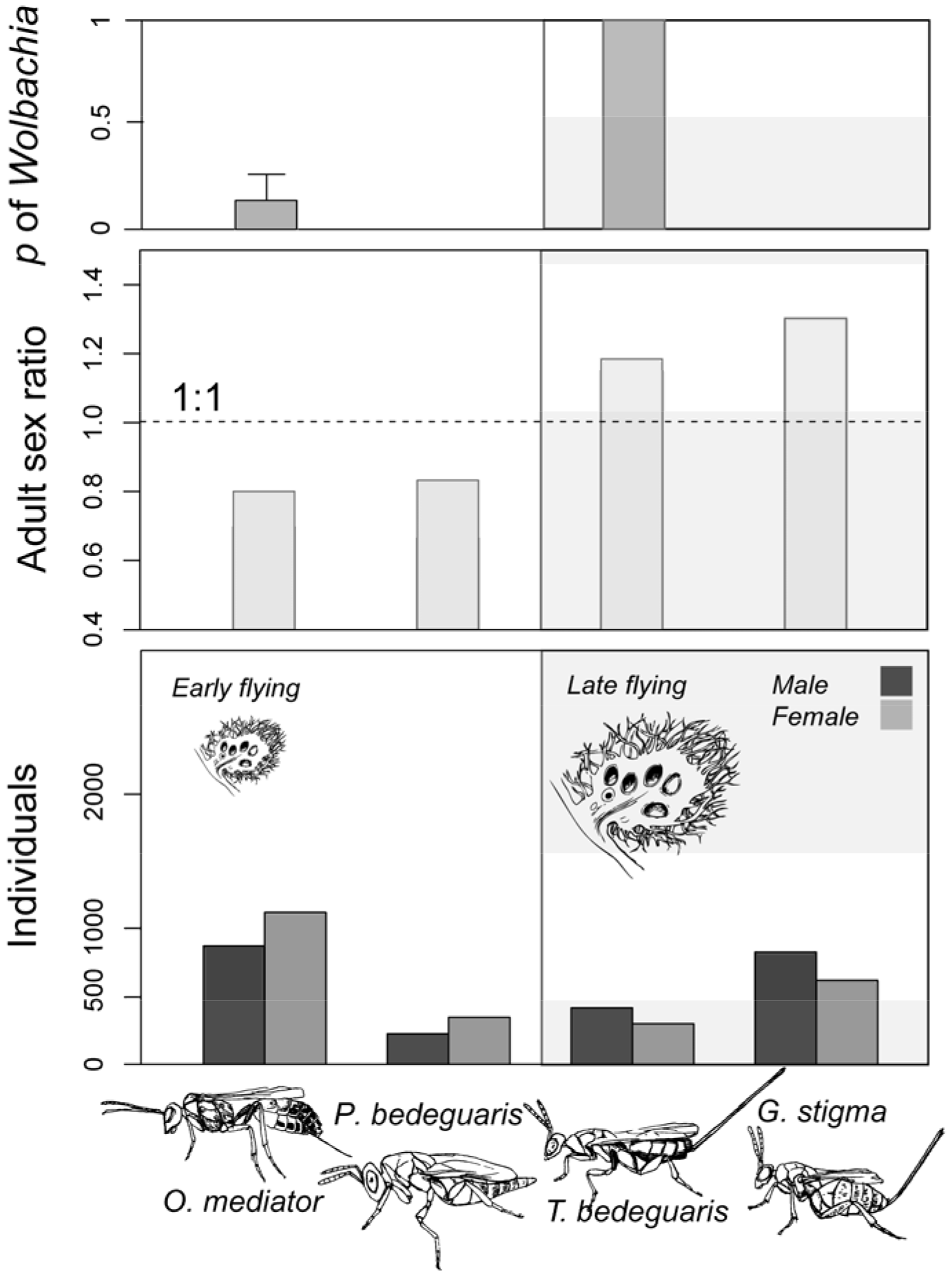
Studied parasitoids. Emereged individual numbers, mean adult sex ratio (ASR) and prevalence of *Wolbachia* infection (mean ± SD) of early and late flying parasitoid species emerged from rose galls (*Diplolepis rosae*).

### Infection of parasitoids by endosymbionts

To evaluate presence of *Wolbachia* endosymbionts, out of N=241 parasitoids, specimen numbers for chosen species were: *O. mediator* N=47, *P. bedeguaris* N=54, *T. bedeguaris* N=69, *G. stigma* N=71. Additionally, presence of *Cardinium* was also tested, by analysing 2 females and 2 males (in total N=16 specimens) of each species. The specimens analysed for *Wolbachia* and *Cardinium* presence were selected individually from different galls collected from different sites, thus no parasitoid specimens were sharing the same gall.

Genomic DNA was extracted using a commercial kit (ISOLATE II Genomic DNA Kit, Bioline) following the manufacturer’s protocol, and was checked for wasp DNA by PCR amplification of a mitochondrial COI sequences (LCO1490/HCO2198 primer pair (Folmer, Black, Hoeh, Lutz, & Vrijenhoek, 1994)). PCR products were purified with a commercial kit (Promega, Wizard SV Gel and PCR Clean–Up System, USA) and sent for sequencing to Macrogen Inc. (Korea). Sequences were verified using the Basic Local Alignment Search Tool (https://blast.ncbi.nlm.nih.gov/Blast.cgi).

The presence or absence of *Wolbachia* was tested by amplifying the *wsp* gene (81F/691R primer pairs (Braig, Zhou, Dobson, & O’neill, 1998)). PCR was performed in a 25 μl reaction volume at an annealing temperature of 50°C (*D. rosae*) or 42°C (*O. mediator and G. stigma*). PCR reactions were checked with both a positive (known infected individual) and a negative control (water). PCR products were visualized on a 1% agarose gel. For samples were the PCR product was absent, in order to confirm the absence of infection two other *Wolbachia* specific markers, the 16S RNS gene (primer pair 99F/994R (O’Neill, Giordano, Colbert, Karr, & Robertson, 1992)) and the *fstZ* gene (FtsZ-F/FtsZ-R primers (Werren, 1997)), were amplified.

The presence or absence of *Cardinium* was tested with the Ch-F/Ch-R primer pair at 57°C. Primers were designed to identify *Cardinium* and other related *Bacteroidetes* symbionts (Zchori-Fein & Perlman, 2004).

### Landscape characteristics

We analysed agricultural and habitat cover types from landscapes within a radius of 2.5 km of each site’s surroundings (Fig. 1). Maps were clipped from Corine Land Cover (Büttner et al., 2002; CLC, 2006, Version 18.5.1) vector overlays by having centroids the mean centroid of surveyed plots of a given landscape. Around each landscape’s centroid a circular buffer was drawn in Quantum GIS (version 2.14.7 “Essen”; QGIS Development Team, 2016), then vector overlays were intersected with these 5 km diameter circular polygons. Areas, edge lengths, patch numbers of agricultural and habitat cover types within the maps were calculated using the package LecoS (Martin Jung, 2016) in Quantum GIS.

Pastures with rose shrubs and shrub encroached grasslands were the gall inducer’s habitat: within its patches (habitat cover type) abundance of host plants (wild roses, *Rosa* sp.) was greater. We calculated mean patch area (McGarigal & Marks, 1995) of habitat cover type, shape index of agricultural cover type (McGarigal et al., 1995; McGarigal, 2014) and agricultural cover type variability through time (landscape history) within each landscape. Mean patch area equals sum of corresponding patch metric values, divided by number of patches of the same type (McGarigal, 2014). Shape index equals patch perimeter divided by the square root of patch area, adjusted by a constant to adjust for a square standard (0.25) (McGarigal, 2014). Ratio of agricultural cover type for each plot was calculated as the ratio between the total area of agricultural cover type and total area of the plot. Landscape history was calculated as the CV% (coefficient of variation percent) of the analysed plot’s agricultural cover types from three different Corine Land Cover (Büttner et al., 2002) maps for years 1990, 2006 and 2012

### Data analysis

We analysed parasitoids (Table 1) emerged from altogether N=617 *D. rosae* galls collected from N=196 rose shrubs (*Rosa* sp.). Data were analysed in the statistical computing environment R version 3.3.1 (R Development Core Team, 2016). We used nested binomial GLMM’s on N=617 ASR values. We used as outcome variable the ASR of the separated parasitoid species emerging from one gall. The parasitoid species’ ASR was calculated as male to female ratio from each sampling unit (individual gall). We made two set of analyses: one for all parasitoids, i.e. the data set containing all galls, and one for separate parasitoids, in which we took the data sets containing only those galls from which the analysed species emerged. Independent variables on local scale were: (1) parasitoid species phenology: two level factorial variable (early and late flying species), (2) diameter (mean of three perpendicular diameters) and density of galls (per sampling plots). Independent variables on landscape scale were: (1) shape index of agricultural patches, (2) mean habitat patch area and (3) landscape history. Collection years, sites, plots, bushes and species were included into models as nested random effects. Presence and percentage of *Wolbachia* infection were used as random variables. We performed logistic GLMMs with package *lme4* with binomial error distributions (Bates, Mächler, Bolker, & Walker, 2015). Variables had variance inflation factor values smaller than 4, thus collinearity was not an issue (Zuur, Ieno, Walker, Saveliev, & Smith, 2009; Dormann, Elith, Bacher, Buchmann, Carl, et al., 2013).

**Table 1.**
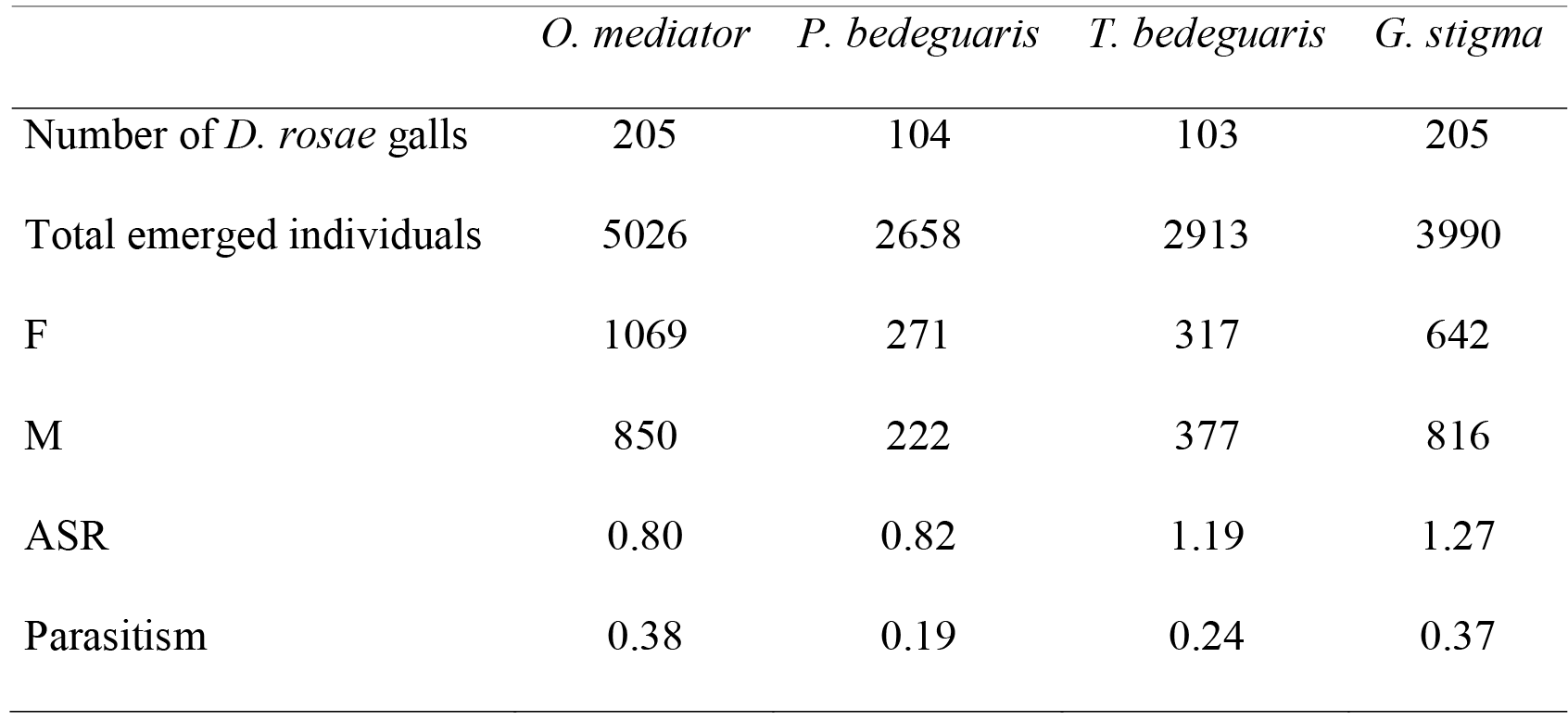
Number of collected galls, total emerged individuals, males and females, adult sex ratio (ASR) and parasitism ratios of the four parasitoid species emerged from N=617 rose galls (*Diplolepis rosae*).

## Results

ASR of *O. mediator* and *P. bedeguaris* was female biased, while of *G. stigma* and *T. bedeguaris* was male biased (Table 1, Supplementary Material Table S1 and Fig. 2). ASR was significantly different between early and late flying species pairs (GLMM: χ^2^=38.06, df=1, p<0.001) and decreased significantly with gall size increase (GLMM: χ^2^=10.17, df=1, p=0.001) (Fig. 3). These biases showed no significant changes between years (GLMM: χ^2^=0.47, df=2, p=0.79) and sites (GLMM: χ^2^=3.71, df=6, p=0.72). Gall numbers decreased with increasing mean habitat patch area (negative binomial GLM: estimate=-0.26, SE=0.03, z= −8.94, p < 0.001), while gall diameter decreased with increasing gall numbers (LM: estimate=−0.62, SE=0.27, t= −2.37, p=0.02) (Fig. 4).

**Fig. 3.**
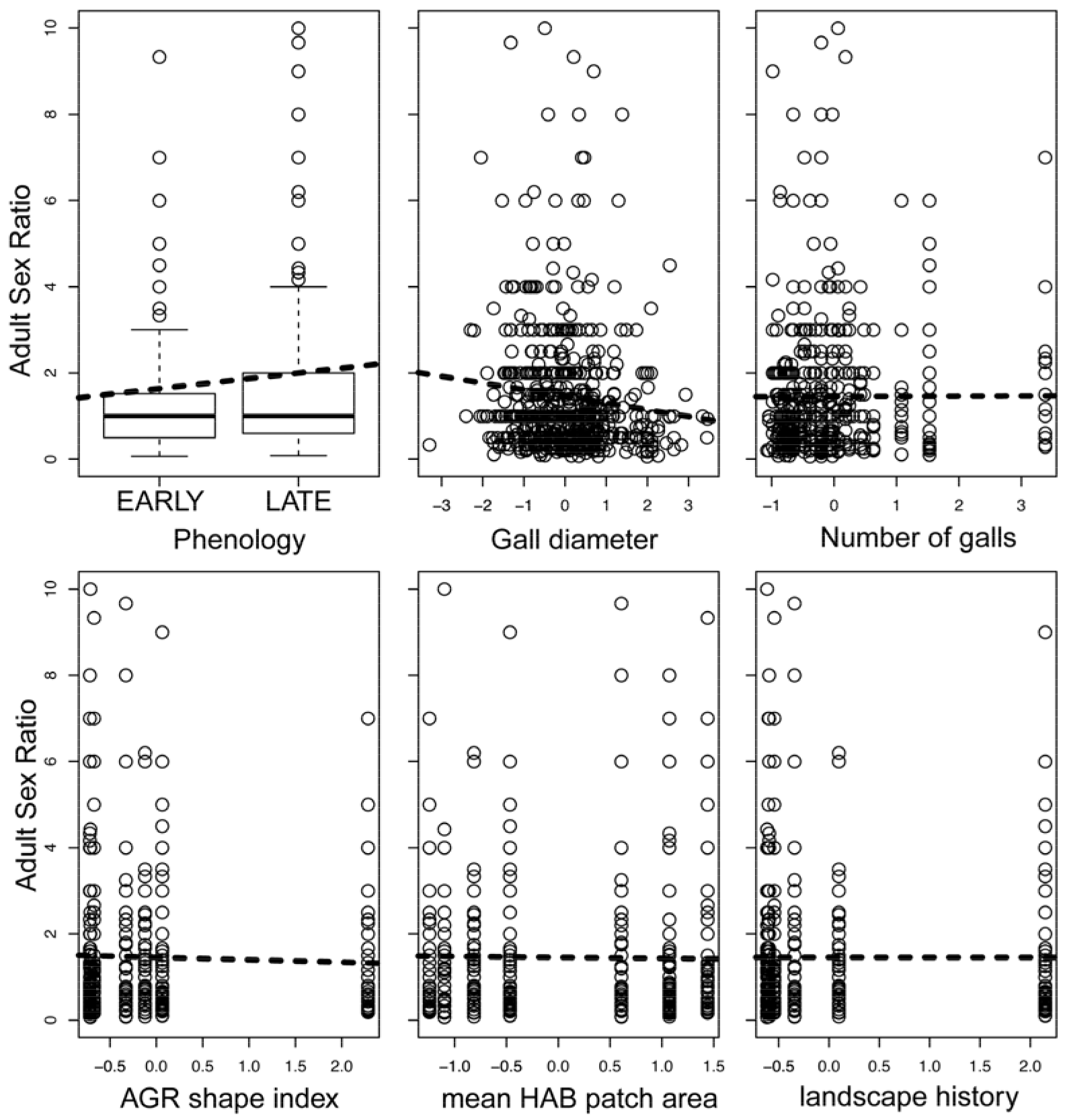
Relationships between the adult sex ratio (ASR) of parasitoids from the community of *Diplolepis rosae* galls and variables as phenology of species and environmental ones on local and landscape scale (logistic mixed effect linear models). Local scale variables: gall diameter and number of galls per bush. Landscape scale variables: shape index of agricultural (AGR) patch types, mean habitat (HAB) patch area and landscape history. All independent variables were scaled.

**Fig. 4.**
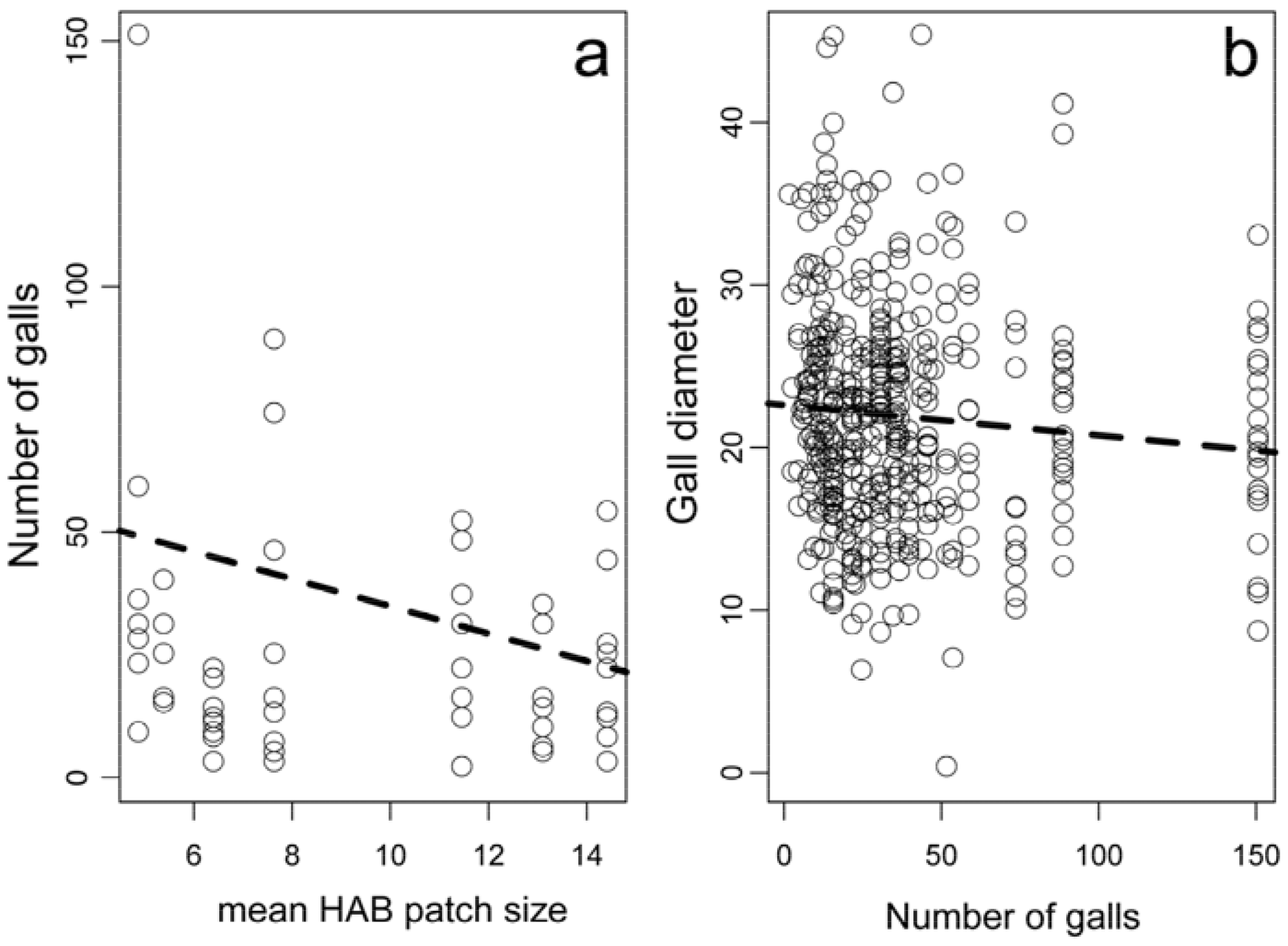
Relationships between small and large scale varaibles: decrease of the number of galls along increasing mean habitat (HAB) patch area and decrease gall diameter along increasing number of galls.

*Wolbachia* was not detected in *P. bedeguaris* and *G. stigma*, but in *O. mediator* and *T. bedeguaris* its presence was confirmed (Fig. 2). Regardless of their sex all *T. bedeguaris* specimens were infected by *Wolbachia*, while in *O. mediator* its prevalence considering both sexes varied around 23.33% (±19.66%) (females: 36.66% (±32.04%), males: 10% (±15.49%)) (Table 2). *Wolbachia* incidence did not affect the parasitoid community’s ASR (GLMM: χ^2^=0.87, df=1, p=0.35), and neither did prevalence (GLMM: χ^2^=3.31, df=1, p=0.07). The other endosymbiont, *Cardinium* was not present in the analysed parasitoids, therefore we didn’t pursue this analysis further.

**Table 2.** Probability of *Wolbachia* infection in analysed samples from the surveyed sites (N=241).

|  | *O. mediator* | *P. bedeguaris* | *T. bedeguaris* | *G. stigma* |
| --- | --- | --- | --- | --- |
| site1 | 0.3 (N=12) | 0 (N=18) | 1 (N=18) | 0 (N=18) |
| site2 | 0.5 (N=6) | na (N=0) | 1 (N=5) | 0 (N=6) |
| site3 | 0.0 (N=5) | na (N=0) | 1 (N=5) | 0 (N=6) |
| site4 | 0.0 (N=4) | 0 (N=6) | 1 (N=6) | 0 (N=6) |
| site5 | 0.3 (N=6) | 0 (N=6) | 1 (N=6) | 0 (N=6) |
| site6 | na (N=0) | 0 (N=6) | 1 (N=6) | 0 (N=6) |
| site7 | 0.3 (N=14) | 0 (N=18) | 1 (N=23) | 0 (N=23) |

Neither small nor large scale variables affected the ASR of the parasitoid community: gall numbers (GLMM: χ^2^=2.49, df=1, p=0.11), bush numbers (GLMM: χ^2^=1.06, df=1, p=0.30), shape index of agricultural patches (GLMM: χ^2^=0.00, df=1, p=0.98), mean habitat patch area (GLMM: χ^2^=0.08, df=1, p=0.77) and landscape history (GLMM: χ^2^=0.28,df=1, p=0.6)(Fig. 2).

The interaction between mean habitat patch area and phenology of parasitoids was associated significantly with parasitoid ASR (Table 3). Parasitoid ASR was associated significantly to parasitoid wasp phenology: parasitoid ASR was larger for late flying species than for early flying ones. Parasitoid ASR decreased significantly with increasing gall sizes. While landscape scale effects had no significant effects on parasitoid ASR, the interaction of parasitoid phenology with mean habitat patch area was significant. While mean habitat patch area has no effect on early flying species ASR, for late ones ASR decreased with increasing mean habitat patch area (Fig. 5).

**Table 3.** Significance of variables in GLMM’s analysing the adult sex ratio (ASR) of the parasitoid community inhabiting galls of *Diplolepis rosae* (N=617). In the table are presented the slopes, their standard errors, z-and p-values belonging to a single linear model. AGR is the abbreviation of agricultural, while HAB of habitat. ***: p<0.001; *: p<0.5; •: p<1.0.

|  | estim. | SE | z | p |  |
| --- | --- | --- | --- | --- | --- |
| (Intercept) | -0.22 | 0.06 | -3.72 | <0.001 | *** |
| phenology (LATE Vs. EARLY) | 0.41 | 0.07 | 5.52 | <0.001 | *** |
| gall diameter | -0.14 | 0.04 | -3.98 | <0.001 | *** |
| number of galls | 0.09 | 0.06 | 1.63 | 0.103 |  |
| AGR shape index | 0.05 | 0.09 | 0.57 | 0.566 |  |
| mean HAB patch area | 0.12 | 0.09 | 1.41 | 0.159 |  |
| landscape history | 0.03 | 0.06 | 0.55 | 0.584 |  |
| phenology: AGR shape index | -0.20 | 0.11 | -1.79 | 0.073 | • |
| phenology: mean HAB patch area | -0.23 | 0.10 | -2.27 | 0.023 | ** |
| phenology: landscape history | -0.15 | 0.08 | -1.80 | 0.071 | • |

**Fig. 5.**
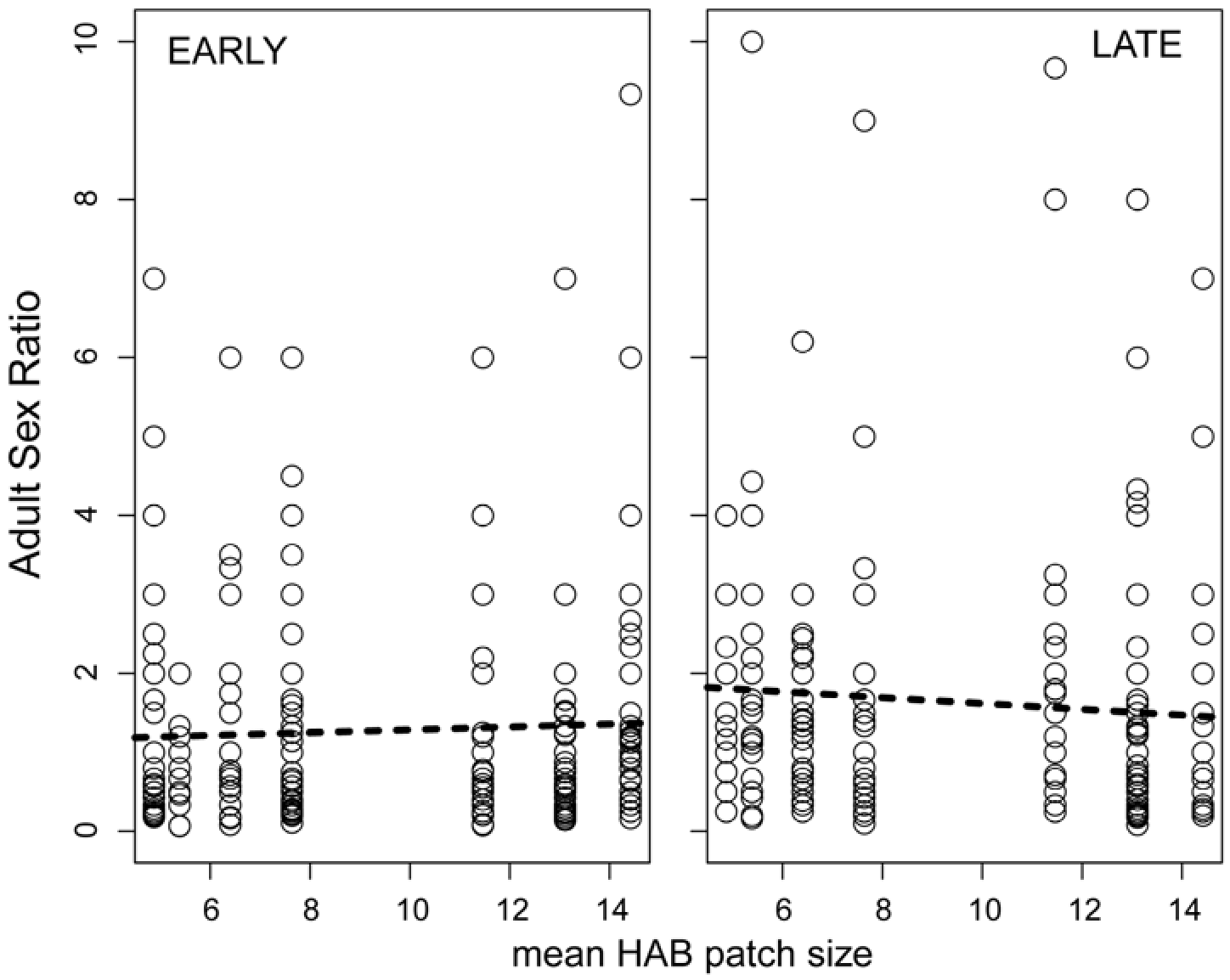
The difference between the relationship of adult sex ratio (ASR) of early (*O. mediator* and *P. bedeguaris*) and late (*G. stigma* and *T. bedeguaris*) flying parasitoids with the mean habitat (HAB) patch size (logistic mixed effect linear model).

When analysing parasitoid species separately we found different variable pattern affecting ASR than that affecting the community pattern (Table 4). *Wolbachia* prevalence in *O. mediator* was the most significant explaining variable of the ASR, and was followed by the gall diameter. With increasing *Wolbachia* prevalence the ASR of *O. mediator* increased significantly. With the increasing gall diameter the ASR of *O. mediator* significantly decreased. For *P. bedeguaris* only gall diameter showed significant effect on ASR (Table 4). With the increasing gall diameter the ASR of *P. bedeguaris* decreased significantly. Not even gall diameter was a significant explaining variable of *T. bedeguaris* ASR. For *G. stigma* gall diameter and mean habitat patch ratio were associated only marginally significantly the ASR (Table 4). With both previously mentioned variables the ASR of G. stigma decreased significantly.

**Table 4.** Significance of variables in GLMM’s analysing the adult sex ratio (ASR) of different parasitoid species belonging to the community of *Diplolepis rosae*. In the table are presented the slopes, their standard errors, z-and p-values belonging to four separate linear models. AGR is the abbreviation of agricultural, while HAB of habitat. **: p<0.01; *: p<0.5;: p<1.0.

|  | estim. | SE | z | p |  |
| --- | --- | --- | --- | --- | --- |
| *ASR of O. mediator* |  |  |  |  |  |
| (Intercept) | -0.52 | 0.17 | -3.05 | 0.002 | ** |
| *Wolbachia* prevalence | 1.20 | 0.52 | 2.31 | 0.021 | * |
| gall diameter | -0.13 | 0.05 | -2.41 | 0.016 | * |
| number of galls | 0.01 | 0.09 | 0.08 | 0.932 |  |
| AGR shape index | -0.06 | 0.14 | -0.44 | 0.662 |  |
| mean HAB patch area | -0.04 | 0.15 | -0.24 | 0.807 |  |
| landscape history | 0.04 | 0.11 | 0.38 | 0.701 |  |
| *ASR of P. bedeguaris* |  |  |  |  |  |
| (Intercept) | -0.19 | 0.09 | -2.02 | 0.043 | * |
| gall diameter | -0.22 | 0.10 | -2.28 | 0.023 | * |
| number of galls | 0.20 | 0.12 | 1.58 | 0.114 |  |
| AGR shape index | 0.01 | 0.16 | 0.06 | 0.948 |  |
| mean HAB patch area | 0.09 | 0.14 | 0.62 | 0.534 |  |
| landscape history | 0.05 | 0.10 | 0.48 | 0.629 |  |
| *ASR of T. bedeguaris* |  |  |  |  |  |
| (Intercept) | 0.18 | 0.09 | 1.96 | 0.050 | • |
| gall diameter | -0.11 | 0.09 | -1.33 | 0.184 |  |
| number of galls | -0.04 | 0.11 | -0.34 | 0.737 |  |
| AGR shape index | 0.17 | 0.16 | 1.03 | 0.303 |  |
| mean HAB patch area | 0.11 | 0.15 | 0.74 | 0.456 |  |
| landscape history | 0.18 | 0.11 | 1.68 | 0.093 | • |
| *ASR of G. stigma* |  |  |  |  |  |
| (Intercept) | 0.22 | 0.09 | 2.41 | 0.016 | * |
| gall diameter | -0.13 | 0.07 | -1.79 | 0.074 | • |
| number of galls | 0.11 | 0.10 | 1.08 | 0.282 |  |
| AGR shape index | -0.15 | 0.11 | -1.39 | 0.166 |  |
| mean HAB patch area | -0.20 | 0.11 | -1.87 | 0.062 | • |
| landscape history | -0.13 | 0.09 | -1.39 | 0.163 |  |

The parasitism of the analysed parasitoid species ranged between 0.19% and 0.38% (Table 1). The overall parasitism of the analysed galls was 49.68%. 65.21% of analysed galls contained only one parasitoid species, while 28.11% contained two parasitoid species. 5.99% of the analysed galls were parasitized by three species and only 0.69% were simultaneously parasitized by all four species. Of the encountered 28.11% binal parasitoid occurences 67.21% were associations of early and late flying species, only 32.79% were associations between species of the same phenology. Mean size of galls with one parasitoid is 20.61 mm, while mean size of those with two parasitoid species is 22.5 mm. Galls with one parasitoid are significantly smaller than those with two parasitoid species (Welch two sample t-test: t=3.38, df=524.21, p=0.0007).

## Discussion

We have found that ASR of parasitoids belonging to the community of Robin’s pincushion gall (*D. rosae*) may depend on host availability through local resource competition (LRC), if the analysed parasitoids exhibit intraspecific competition. At least it depends more on environmental variables than on the presence of internal symbionts. We found that a large scale landscape variable, habitat availability indirectly affects the ASR. We also found species specific responses: *O. mediator* was considerably affected by *Wolbachia*, while late flying species were not affected by either of analysed variables.

### Infection by endosymbionts

*Wolbachia* infection was present in two cases: *O. mediator* was infected with a changing prevalence, while *T. bedeguaris* was uniformly infected. One of them is an early, the other is a late flying species, which means that *Wolbachia* infection has no effect on the parasitoid community’s ASR pattern.

For *O. mediator* a study found 10% prevalence of *Wolbachia* (Kohnen, Richter, & Brandl, 2012), while another showed that out of three analysed localities, only specimens from one were infected by *Wolbachia* (Schilthuizen & Stouthamer, 1998). Thus, it seems that although *O. mediator* is infected, prevalence of *Wolbachia* is low. *T. bedeguaris* in the first study (Kohnen et al., 2012) was not a target species, while in the second (Schilthuizen et al., 1998) all its specimens were infected. Our results are in concordance with the previous studies since all *T. bedeguaris* specimens were infected by *Wolbachia* also in our samples.

*G. stigma* showed in both studies (Schilthuizen et al., 1998; Kohnen et al., 2012) a complete lack of *Wolbachia*, as it did in our samples. The only difference between our results and literature concerns the species *P. bedeguaris*: in a study 6 specimens were analysed and they found the Type I strain of *Wolbachia* (Schilthuizen et al., 1998), while we did not found any evidence for *Wolbachia* presence (N=36). There are two possibile explanations for this difference: *Wolbachia* is missing from the eastern Carpathian Basin from *P. bedeguaris*, or it is present but we have not detected it. The first possibility is more likely to be true since, after checking for the *wsp* gene, all negative results were further proofed with two additional *Wolbachia* specific markers (16S RNS gene of the *Wolbachia* and the *fstZ*).

Infection pattern of *Wolbachia* in the eastern Carpathian Basin, excepting *P. bedeguaris*, resembles the European pattern (Schilthuizen et al., 1998; Kohnen et al., 2012). *Wolbachia* effects on reproduction pattern of the studied parasitoids are not known, therefore much about *Wolbachia* impact on the ASR cannot be said (Cook & Butcher, 1999). The only species where *Wolbachia* infection alters the ASR is *O. mediator* where prevalence varies significantly. One mechanism of *Wolbachia* to affect their hosts is CI. Often this has weak effects (few affected progeny) and thus influences only slightly their progeny’s ASR.

Our results showed no *Cardinium* infection, which means that even if it is present in may have a low prevalence. *Cardinium* is rarer in insects than *Wolbachia* (Floate et al., 2013). In the Chalcidoidea superfamily, *Cardinium* has been found only in Aphelinidae, Encyrtidae and Eulophidae. Species belonging to Torymidae have not been analysed, while one species belonging to Pteromalidae showed no *Cardinium* presence. In the Ichneumonidae family only one species was analysed but it was also lacking *Cardinium* (Zchori-Fein et al., 2004).

### Phenology and functional trait

We have found that parasitoid phenology, which correlates with a functional trait: the ovipositor sheath length, is strongly associated with parasitoid ASR’s variability. This ASR difference is due to the fact that early flying species exhibited female biased, while late flying species exhibited male biased sex ratios. Stille (Stille, 1984) reported for *O. mediator* an ASR of 0.612, while for *T. bedeguaris* of 1.104. This pattern coincides with our findings and means that early flying species have smaller ASR at a large (at least European) scale. But which variables cause the ASR difference between the early and late flying species?

Firstly, parasitoid ASR is affected through their phenological sequence. Early flying species actually find all larvae parasitized, while late flying species find a lot of already parasitized larvae by early species. Therefore, late flying species may face a higher LRC, which leads to higher male production, and thus higher ASR (West, 2009).

Secondly, late flying species’ encounter larger gall chambers and thicker chamber walls compared to early flying species. Thus, this difference means that late species flying in the summer have to overcome also increased gall chambers and diameters. Thus, late flying species resource availability is decreased manifold, also by the fact that spherical galls with larger diameters have less chambers on their surfaces than in their inside, and so in spite of their longer ovipositors these species can reach to less host larvae than the early ones can (László & Tóthmérész, 2013). Differing ovipositor sheath lengths between early and late flying species shows that late flying species are adapted morphologically to LRC. LRC thus may also affect these species ASR.

### Local variables: gall diameter

The second most significant variable affecting parasitoid ASR was host availability through gall diameter. As gall diameter increased, ASR decreased for all four species. In galls with large diameters gall chamber diameters are also larger than in small galls (László et al., 2013). Thus, in large galls there will presumably be larger gall inducer larvae than in small galls. As parasitoid females produce daughters where large host larvae are present (Charnov et al., 1981), when female parasitoids find large galls with large larvae, they will lay eggs from which daughters will develop, and so ASR decreases. This relationship may affect the female bias found in early species, but cannot overcome the LRC affecting late flying species, because gall diameter affects less ASR than phenology does.

### Landscape variables: mean habitat patch area

The variable for which the relationship with parasitoid ASR showed significant difference between early and late flying species belonged to large scale variables (Fig. 5). Late flying species *G. stigma* and *T. bedeguaris* showed high ASR and their ASR showed an increasing trend towards small mean habitat patch area.

The mechanism through which mean habitat patch area may affect the ASR of late flying parasitoids may be the following: i) in small habitats gall number is higher than in large ones (Fig. 4a), ii) at high gall number, gall diameter is smaller than at small gall number (Fig. 4b), iii) small galls exhibit high ASR (Fig. 3, upper middle) due to LRC. It is known that parasitoid ASR is affected by several variables (King, 1987; Fox, Letourneau, Eisenbach, & Nouhuys, 1990), but these are mostly local. Large scale variables were scarcely analysed, but based on our results we can assume they have an effect on the ASR of parasitoids, even if only indirectly.

Habitat size may be small due to habitat loss and fragmentation (Fahrig, 2003) and may increase isolation by distance (Amos et al., 2014). Our knowledge on how fragmentation affects ASR is limited and comes largely from vertebrate systems (Harrisson, Pavlova, Amos, Radford, & Sunnucks, 2014; Reid et al., 2014), but insects with short generation times present an ideal opportunity to study these questions (Murphy, Battocletti, Tinghitella, Wimp, & Ries, 2016).

### *Wolbachia* and other variable effects on *O. mediator* ASR

The only parasitoid for which *Wolbachia* incidence explained significantly the ASR was *O. mediator*. Even though there is no information regarding *Wolbachia*’s effect mechanism on this species (Schilthuizen et al., 1998; Kohnen et al., 2011), it is known that *Wolbachia* presence usually is linked to small ASRs; in this way the endosymbiont’s high inheritance is assured (Charlat, Hurst, & Merçot, 2003; Werren, Baldo, & Clark, 2008). For *O. mediator* the small ASR may indicate such mechanism; however, resource availability effect remains also important (Table 4). Thus, internal and local environmental variables affect together *O. mediator*’s ASR. Studies that target parasitoids regarding internal and local variables in relation with ASR are rarely reported (Duplouy et al., 2015). In one case, spite of *Wolbachia* infection the ASR was distorted the same way as in uninfected individuals (Abe, Kamimura, Kondo, & Shimada, 2003). Our point is that *Wolbachia* infection may have great impact on their hosts ASR, but besides endosymbionts, other environmental variables as host availability are also highly important.

### Conclusions

We have found significantly biased ASRs in a parasitoid community belonging to the same host: the bedeguar gall. Phenology of species explained the variability of ASR the most. Phenology is linked to functional traits such as ovipositor length and environmental variables as host availability through gall size and competition. These variables have shown to be more important than the presence of the endosymbiont *Wolbachia* for three of the four analysed species. Moreover, we found an indirect effect on the parasitoid community’s ASR of a large scale variable, the mean habitat patch size. We conclude that large scale effects are also important in shaping the parasitoid ASR.

## Acknowledgements

Molecular processing for all specimens was done at the Interdisciplinary Research Institute on Bio–Nano–Sciences of BBU, Cluj. We thank for the help of K. Sólyom, Á. Lubinsky, H. Prázsmári and T. I. Kelemen in specimen identifications and selections during the preparation for DNA extraction. The work of ZL was supported by a grant of the Romanian Ministry of Education, CNCS – UEFISCDI, project number PN-II-RU-PD-2012-3-0065 and by an internal grant of UBB, Cluj-Napoca, with project number BBU-GTC-2016-31796. The work of BT was supported by TÁMOP-4.2.2.B-15/1/KONV-2015-0001. During the preparation of the manuscript, ALD received financial support from the Collegium Talentum scholarships, Hungary.

Author contributions. ZL initiated the project, made landscape maps and species determinations. ALD and LK made molecular analyses. ZL analysed, interpreted data and drafted the manuscript. BT contributed substantially to revisions. All authors gave final approval for publication.

